# *Cassava brown streak virus* has a rapidly evolving genome: implications for virus speciation, variability, diagnosis and host resistance

**DOI:** 10.1101/053546

**Authors:** Titus Alicai, Joseph Ndunguru, Peter Sseruwagi, Fred Tairo, Geoffrey Okao-Okuja, Resty Nanvubya, Lilliane Kiiza, Laura Kubatko, Monica A. Kehoe, Laura M. Boykin

## Abstract

Cassava is a major staple food for about 800 million people in the tropics and subGtropical regions of the world. Production of cassava is significantly hampered by cassava brown streak disease (CBSD), which is caused by *Cassava brown streak virus* (CBSV) and *Ugandan cassava brown streak virus* (UCBSV). The disease is suppressing cassava yields in eastern Africa at an alarming rate. Previous studies have documented that CBSV is more devastating than UCBSV because it more readily infects both susceptible and tolerant cassava cultivars, resulting in greater yield losses. Using whole genome sequences from NGS data, we produced the first coalescentGbased species tree estimate for CBSV and UCBSV. This species framework led to the finding that CBSV has a faster rate of evolution when compared with UCBSV. Furthermore, we have discovered that in CBSV, nonsynonymous substitutions are more predominant than synonymous substitution and occur across the entire genome. All comparative analyses between CBSV and UCBSV presented here suggest that CBSV may be outsmarting the cassava immune system, thus making it more devastating and harder to control.

## Introduction

Cassava (*Manihot esculenta* Crantz) is a major staple food crop for over 800 million people in over 100 tropical and subGtropical countries^1^. In subGSaharan Africa, it is the main source of dietary calories for approximately 300 million people^2^. The tuberous storage roots of cassava are rich in carbohydrates and can be cooked or processed for human food, animal feeds and a wide range of industrial products. The crop is relatively drought tolerant and can yield well even in less fertile soils, hence, its importance to poor families farming marginal lands^3^. Cultivation of cassava is most adversely affected by two viral diseases: cassava mosaic disease (CMD) and cassava brown streak disease (CBSD)^4^, which together were reported to cause production losses of more than US$1 billion every year^5^ in Africa.

Serious yield losses due to CMD were first observed on mainland East Africa in the 1920s^6^. Recorded epidemics of CMD later occurred in the 1930s, 1940s and from 1990s to date^7^,^8^. By contrast, for about 70 years since it was first described^9^, CBSD was confined to low altitudes (below 1000 meters above sea level) along coastal eastern Africa in Kenya, Tanzania, Mozambique and Malawi. However, in the early 2000s, outbreaks of CBSD were reported over 1000 km inland at midGaltitude locations (above 1000m) in multiple countries all around Lake Victoria in Uganda^10^, western Kenya^11^ and northern Tanzania^4^. Where it is already established in eastern Africa, the current CBSD epidemic prevails as the main cause of losses in cassava production. Over the last 10 years, the CBSD epidemic has expanded to other countries in East and Central Africa such as Rwanda, Burundi, Congo, DR Congo and South Sudan^12^-^14^. This has significantly increased the risk to countries in central and west Africa which are among the world’s leading cassava producers, and where CBSD does not occur.

CBSD is caused by *Cassava brown streak virus* (CBSV) and *Ugandan cassava brown streak virus* (UCBSV). Both viruses are (+) ssRNA viruses in the genus *Ipomovirus* and family *Potyviridae*^15,16,17,18^, and are often together referred to as cassava brown streak viruses (CBSVs). The CBSVs have genomic organization of 10 segments, total size approximately 8.9 to 10.8 kb, and coding for a polypeptide with about 2,900 amino acid residues^15^,^17^,^18^. The complete genome of a CBSD causal virus was first sequenced in 200918, and to date there are only 26 publicly available^19^. Currently there are two species recognized by the ICTV, but Ndunguru et al.^19^ have suggested further speciation in the UCBSV clade. Both viruses are transmitted in a semiGpersistent manner by the whitefly *Bemisia tabaci*^20^ and mechanically^21^. Symptoms of CBSD on cassava vary with cultivar, virus or plant age, but typically include leaf veinal choloris, brown stem lesions, as well as constrictions, fissures and necrosis of the tuberous storage roots^22^,^23^. Overall, both CBSV and UCBSV cause similar symptom types, however, infection with CBSV tend to result in more severe sumptoms.

Although CBSD has become established in eastern Africa, there is limited knowledge on the diversity of causal viruses, their distribution and evolutionary potential. Therefore, it is necessary to obtain several full genome sequences of CBSD viral isolates, better understand the causal viruses and design longGterm control approaches for the disease.

In contrast to the growing knowledge on the causal agents of CBSD, hostGpathogen interactions are less clear. As such, little is known about specific responses of different cassava varieties to prevailing species or strains of CBSD viral pathogens. Development and dissemination of CBSDG tolerant varieties has been the main means adopted for CBSD control in eastern Africa. With significant efforts geared at breeding for CBSDGresistant varieties, it is of great interest to know if such resistance protects cassava against one or both CBSVs. The resistance may be expressed as several related features including restricted infection, systemic spread or recovery of infected plants from disease and the possibility that stem cuttings taken from these may give rise to progeny that are virusGfree (reversion). Recent studies have shown CBSV to be the more aggressive virus, infecting both tolerant and susceptible cultivars as single or mixed infections with UCBSV^15^,^24^,^25^. In contrast, tolerant varieties were infected with only CBSV, but free of UCBSV, suggesting their resistance to the latter. Compared with UCBSV, CBSV isolates have been reported to be more detectable, having higher infection rates by graft inoculation and inducing more severe symptoms^26^. It has also been shown that plants of CBSD tolerant or resistant cultivars graftGinoculated with UCBSV developed milder symptoms and a significantly higher proportion of the progenies were virusGfree (reverted) compared to those infected with CBSV^27^. To date, the underlying reasons for this more aggressive nature of CBSV compared with UCBSV are not known.

In this study, CBSV and UCBSV molecular diversity was investigated by using next generation sequencing to understand new complete genomes of three isolates from Uganda. The sequences obtained were analyzed to determine species composition, CBSV and UCBSV evolutionary rates, potential role of such changes in virusGhost interactions, resulting into cassava cultivar susceptibility or resistance. We set out to answer the following questions:

1. How do the three new complete genomes from Uganda compare to those already published^19^?
2. Are CBSV and UCBSV distinct species and is there further speciation?
3. Why is CBSV more aggressive and harder to breed resistance for than UCBSV?

## Results

### CBSD Field Symptoms Associated with CBSV and UCBSV Isolates

Categorisation of CBSD foliar symptom distribution on symptomatic plants assessed revealed that the most frequently encountered type was LL G symptoms only on lower leaves (68.4%), followed by SW G systemic and on the whole plant (26.3%), and SL – systemic but localized (5.3%) (table 1). Based on CBSVs detected and CBSD leaf symptom severity scores for 57 sampled plants, whereas the majority of plants infected by UCBSV alone as determined by RTGPCR had mild chlorosis (severity score 2), CBSV infections (single or mixture with UCBSV) tended to have moderate to severe symptoms (scores 3G4) in same proportion to those exhibiting score 2 (Fig. 1, Table 1). Regarding the three isolates used here for whole genome sequencing, U8 (UCBSV) was from a plant with CBSD score 3 and LL symptom type. Both CBSV isolates (U1 and U4) were from plants with severity scores 2 and 3, symptom types LL and SL, respectively.

**Figure 1:**
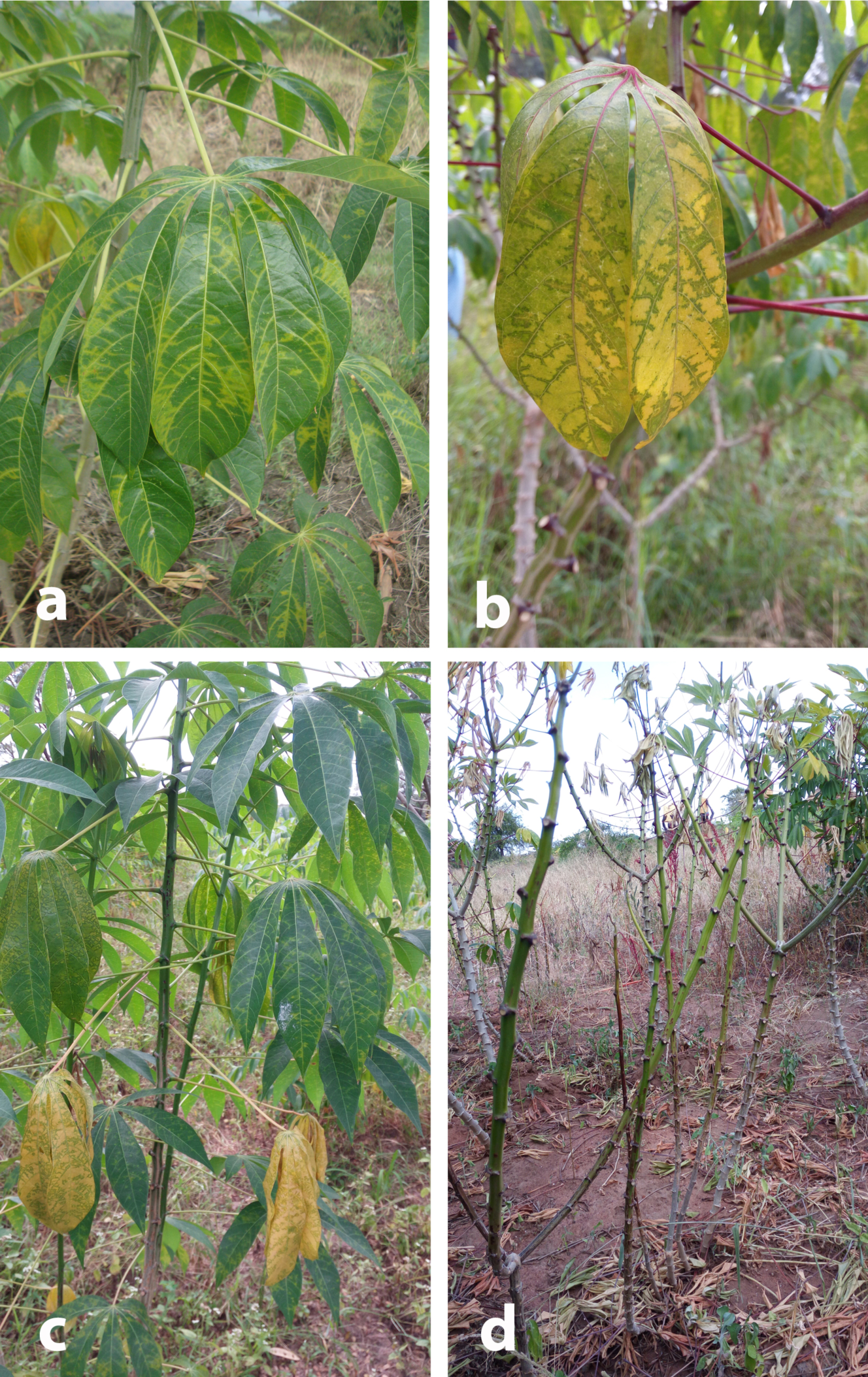
Cassava brown streak disease symptoms on leaves and stems of sampled plants; **(a)** Chlorosis along secondary and tertiary leaf veins of CBSVFinfected plant of cultivar TME 204 (severity score 3), **(b)** Cultivar TME 14 plant with dual CBSV+UCBSV infection showing chlorosis on secondary or tertiary veins, reverse chlorosis (general chlorosis and green area along veins) (severity score 3), **(c)** UCBSVFinfected plant of cultivar TME 204 exhibiting chlorosis on secondary veins, reverse chlorosis, chlorotic spots and mild stem lesions (severity score 3), **(d)** Very severely diseased plant (severity score 5) of cultivar TME 14 infected with both CBSV and UCBSV, and having chlorosis on leaves, severe stem lesions/brown streaks, defoliation, stem dieback.

**Table 1:**
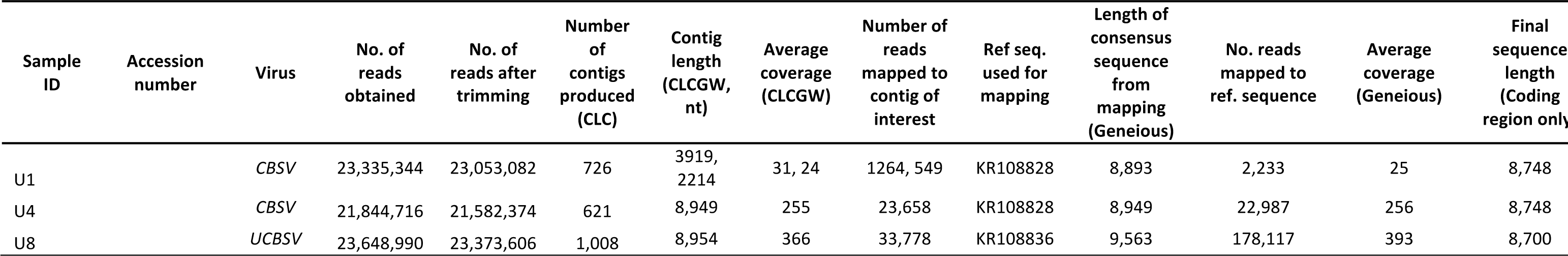
CBSD leaf symptom severities and types on plants infected by *Cassava brown streak virus* and *Ugandan cassava brown streak virus*

1 1Foliar CBSD symptom severity score based on 1-5 scale; 1 = no visible symptoms, 2 = mild vein yellowing or chlorotic blotches on some leaves, 3 = pronounced/extensive vein yellowing or chlorotic blotches on leaves, but no lesions or streaks on stems, 4 = pronounced/extensive vein yellowing or chlorotic blotches on leaves and mild lesions or streaks on stems, 5 = pronounced/extensive vein yellowing or chlorotic blotches on leaves and severe lesions or streaks on stems, defoliation and dieback.

2 Types of foliar CBSD symptoms based on distribution of leaf chlorosis and stem lesions on the plant; systemic and on the whole plant (SW), systemic on leaf or stem parts but localized (SL), only on lower leaves (LL).

### Next Generation Sequencing

The three samples from Uganda produced raw reads ranging from 21,844,716 to 23,648,990. After trimming for quality using CLCGW, these numbers were reduced to 21,582,374 to 23,373,606 (table 2). Following *de novo* assembly of the trimmed reads using CLCGW, the numbers of contigs produced were 621G1,008. The contigs of interest from *de novo* assembly were of lengths 2,214 to 8,954nt, with average coverage 24 to 366. After mapping to a reference genome in Geneious, the lengths of the consensus sequences were 8,893 to 9,563 with average coverages of 25 to 393. The final sequences consisted of a consensus between the *de novo* and the mapped consensus with lengths of 8,700 to 8,748.

**Table 2.**
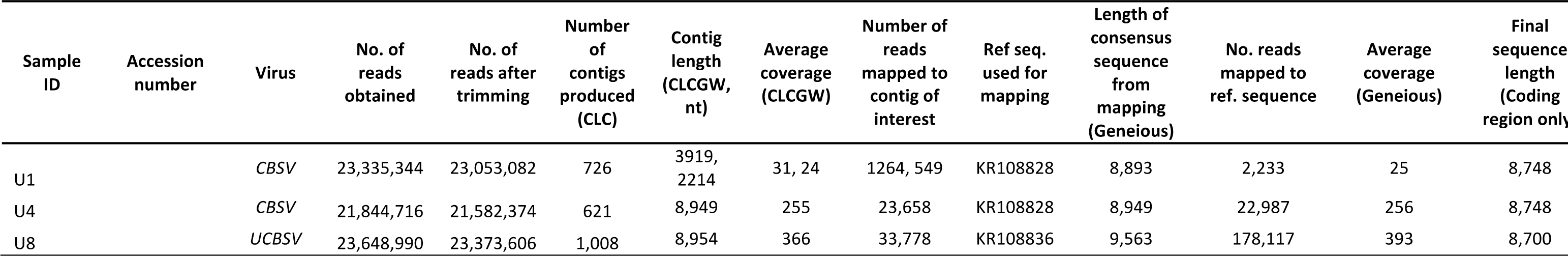
Next generation sequencing data for samples from cassava brown streak disease symptomatic plants collected in Uganda.

### Genomic Variability and Positive Selection

The CBSV genomes included in this study were more variable when compared with those of UCBSV (supplementary figs. S1 and S2). Characterizing amino acid usage at each position in the whole genome revealed that CBSV genomes have nonGsynonymous substitutions present across their entire genome (Fig. 2), and predominating when compared to synonymous substitutions. In contrast, UCBSV had near equal nonGsynonymous and synonymous substitution rates across the entire genome. Genes in the UCBSV genomes with nonGsynonymous substitutions at a higher frequency were: P1, NIb and HAM1 (Fig. 2).

**Figure 2.**
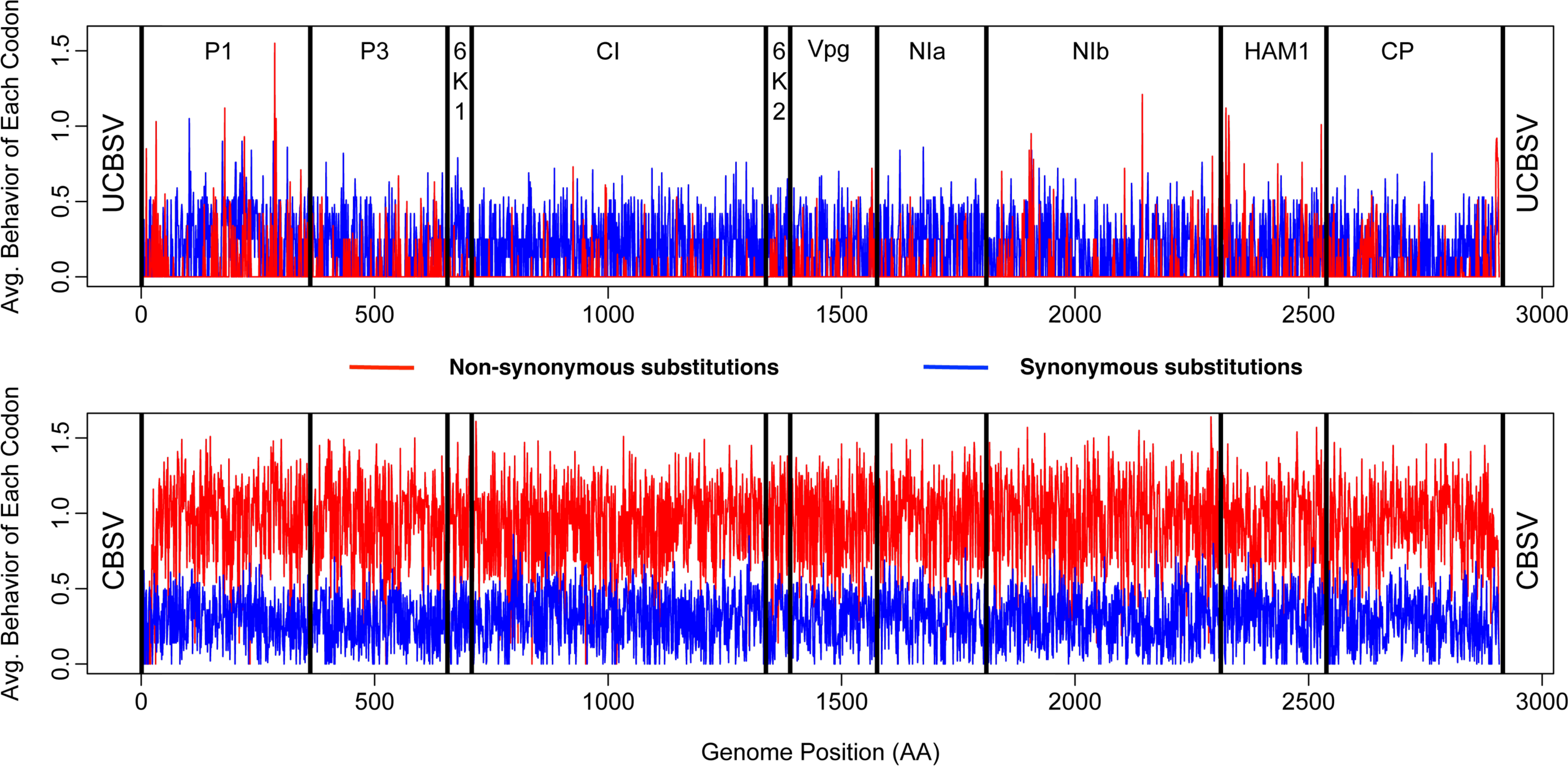
Genetic diversity of CBSV and UCBSV using the <u>S</u>ynonymous <u>N</u>onFsynonymous <u>A</u>nalysis <u>P</u>rogram (SNAP v2.1.1) implemented in the Los Alamos National Laboratory HIVF sequence database (<u>http://www.hiv.lanl.gov</u>)^50^. UCBSV is on the top panel, CBSV at the bottom. The 10 gene segments are labeled from P1FCP.

CBSV had 68 positively selected sites and 66 negatively selected sites, while UCBSV had zero positively selected sites and 558 negatively selected sites (Table 3). Analyzed together there are 3 positively selected sites and 1383 negatively selected sites. The coat protein (CP) of CBSV had the highest number of positively selected sites (16) while 6K2 had zero.

**Table 3.** *Cassava brown streak virus* (CBSV) amino acid (AA) sites under positive selection (analyses method: SLAC HyCPhy). There were no sites under positive selection for *Ugandan cassava brown streak virus* (UCBSV).

| Gene | CBSV AA site under positive selection |
| --- | --- |
| P1 | 44, 46, 50, 174, 224, 230, 250, 283, 288, 349, 358 |
| P3 | 415, 455, 467, 472, 499, 525, 618 |
| 6K1 | 658, 678 |
| CI | 735, 761, 820, 827, 848, 852, 894, 935, 1218 |
| VPg | 1465 |
| NIa | 1620, 1645, 1704, 1754, 1785 |
| Nib | 1879, 1880, 1890, 1907, 1929, 2109, 2145, 2156, 2161, 2285 |
| HAM1 | 2320, 2345, 2404, 2432, 2453, 2475, 2519 |
| CP | 2550, 2555, 2588, 2611, 2631, 2635, 2640, 2659, 2728, 2745, 2783, 2818, 2843, 2860, 2877, 2884 |

### Rates of Evolution

CBSV and UCBSV have different rates of evolution (Table 4). We tested two hypothesis using CODEML. The null hypothesis tested was that CBSV and UCBSV have equal rates of evolution while the alternative hypothesis was that CBSV and UCBSV have different rates of evolution (two omegas; model = 2). The Likelihood Ratio Test was used to test for significance. If the difference in likelihood was greater than 3.84 (based on the ChiGsquared distribution and one degree of freedom) we rejected the null hypothesis that the rates between CBSV and UCBSV are equal.

**Table 4.** Rates of evolution tested using CODEML implemented in PAML. HO was CBSV and UCBSV have equal rates of evolution (one omega; model = 0), while H1 was that CBSV and UCBSV have different rates of evolution (two omegas; model = 2).

| Gene | Assumptions | K (ts/tv rate ratio) | w (omega Dn/Ds) 0 | w (omega Dn/Ds)1 | Likelihood Ratio Test Statistic (if greater than 3.84 reject $H_0$)<br>$H_0$=equal rates |
| --- | --- | --- | --- | --- | --- |
| WGS | UCBSV and CBSV equal rates | 5.90944 | 0.06358 |  |  |
|  | UCBSV and CBSV different rates | 5.9598 | 0.05518 | **0.07622** | 26.29* |
| P1 | UCBSV and CBSV equal rates | 5.07336 | 0.10394 |  |  |
|  | UCBSV and CBSV different rates | 5.05456 | 0.09047 | **0.12203** | 4.61* |
| P3 | UCBSV and CBSV equal rates | 5.17197 | 0.08635 |  |  |
|  | UCBSV and CBSV different rates | 5.20559 | 0.0764 | **0.10198** | 2.22 |
| 6K1 | UCBSV and CBSV equal rates | 19.54143 | 0.00969 |  |  |
|  | UCBSV and CBSV different rates | 19.69676 | **0.01527** | 0.00316 | 3.01 |
| CI | UCBSV and CBSV equal rates | 9.9388 | 0.01722 |  |  |
|  | UCBSV and CBSV different rates | 8.06276 | 0.0155 | **0.01977** | 0.73 |
| 6K2 | UCBSV and CBSV equal rates | 8.40649 | 0.04684 |  |  |
|  | UCBSV and CBSV different rates | 8.82738 | 0.02354 | **0.11057** | 6.74* |
| VPg | UCBSV and CBSV equal rates | 5.852 | 0.05759 |  |  |
|  | UCBSV and CBSV different rates | 5.87009 | 0.054 | **0.06323** | 0.29 |
| NIa | UCBSV and CBSV equal rates | 8.01105 | 0.02932 |  |  |
|  | UCBSV and CBSV different rates | 8.68283 | 0.01408 | **0.06719** | 29.95* |
| NIb | UCBSV and CBSV equal rates | 6.07872 | 0.05329 |  |  |
|  | UCBSV and CBSV different rates | 6.14047 | 0.0452 | **0.06508** | 5.18* |
| HAM1 | UCBSV and CBSV equal rates | 7.25177 | 0.16144 |  |  |
|  | UCBSV and CBSV different rates | 7.2343 | **0.17929** | 0.14007 | 2.23 |
| CP | UCBSV and CBSV equal rates | 13.09297 | 0.06075 |  |  |
|  | UCBSV and CBSV different rates | 13.26752 | 0.05606 | **0.07155** | 1.29 |
|  |  |  | **faster rate UCBSV** | **faster rate CBSV** | * Rates are different |

CBSV whole genome sequences showed it is evolving 5 times faster than UCBSV overall. The genes contributing to this accelerated rate of evolution for CBSV are Nla (D=29.95), followed by 6K2 (D=6.74), Nlb (D=5.18) and P1 (4.61) (Table 4). The transition/transversion ratios were also estimated using CODEML, and show that the 6K1 (19.6) and CP (13.2) genes have the highest estimates while the remaining 8 genes ranged from 5.05 – 9.93.

**Table 5.** Support for Clades A – G (Figure 3) in individual gene trees and whole genome analyses. Table entries represent posterior probabilities from analysis with MrBayes, except values reported for SVDQ, which are bootstrap proportions. Support values below 95% are indicated in bold, and ‘PP’ indicates that the clade was not present.

| Genomic region | Clade A | Clade B | Clade C | Clade D | Clade E | Clade F | Clade G |
| --- | --- | --- | --- | --- | --- | --- | --- |
| P1 | 99.98 | 99.98 | 96.98 | 99.72 | 99.54 | -- | 99.98 |
| P3 | 99.98 | 99.98 | 98.57 | 99.98 | 99.98 | -- | 99.98 |
| 6K1 | **94.66** | 99.85 | 98.87 | 99.95 | -- | -- | 98.42 |
| CI | 99.98 | 99.96 | 99.76 | 99.98 | 99.98 | 99.99 | 99.99 |
| 6K2 | 97.65 | 99.98 | **71.52** | **91.70** | **66.46** | -- | 98.75 |
| Vpg | 99.98 | 99.99 | -- | 99.18 | **64.55** | -- | 99.99 |
| NIa | 99.96 | 99.99 | 99.98 | 99.41 | **76.31** | -- | **89.34** |
| NIb | 99.99 | 99.98 | 99.98 | 99.99 | 99.98 | -- | 99.98 |
| HAM | -- | -- | -- | 99.89 | 99.83 | -- | 98.55 |
| CP | -- | -- | -- | 99.59 | 99.50 | -- | 99.89 |
| Whole genome | 1.00 | 1.00 | 1.00 | 1.00 | 1.00 | 1.00 | 1.00 |
| SVDQ | 1.00 | 1.00 | 1.00 | 1.00 | **87.00** | **44.00** | 1.00 |

### Species Tree Estimation K SVDQ

The species phylogeny (Fig. 3) shows strong support for a split into two primary viral clades, one consisting of CBSV (Fig. 3 clades A and B) and the other consisting of UCBSV (Fig. 3 clades EGG), with 100% bootstrap support separating the two clades. Figure 3 shows clades labeled A–G which correspond to; 1) labels AGF from Ndunguru et al^19^, and 2) a new clade G defined in this study. Within the CBSV clade, there are several additional clades with 100% bootstrap support, including the two new CBSV whole genomes from Uganda (U1 and U4). These are the first CBSV whole genomes sequences from Uganda. The other CBSV grouping with 100% bootstrap support labeled B in Figure 3 contains 4 Tanzania samples KoR6, Tan 79, Tan 19 1 and Nal 07. In the UCBSV clade there are 6 nodes supported with a 100% bootstrap, including the new UCBSV whole genome added from this study (U8) which is sister to Kab 07 from Uganda. In addition, the CBSV clade had all samples from a given country grouping together while the UCBSV clade had monophyletic clades from different countries (the multiGcolored lines in Fig. 3).

**Figure 3.**
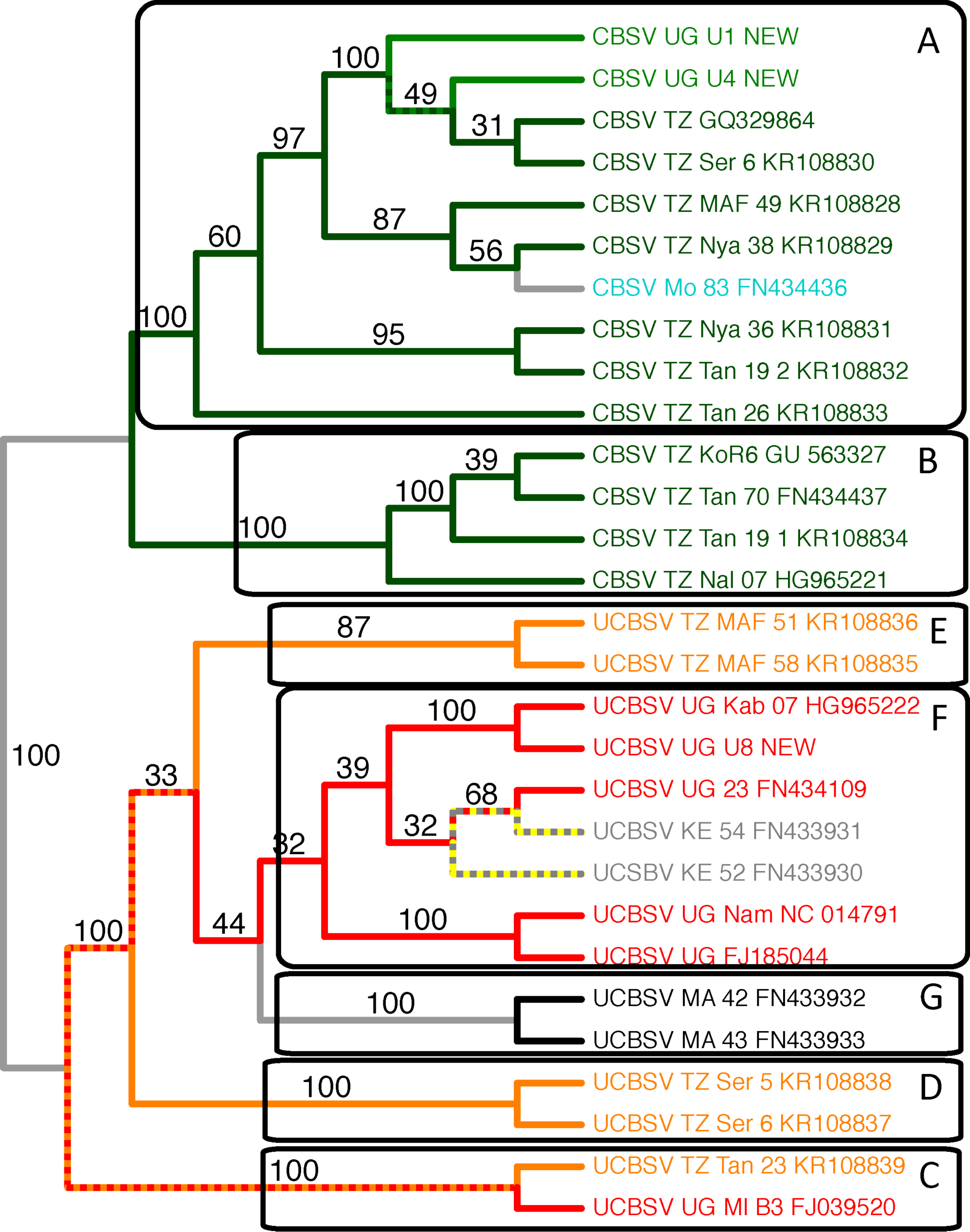
Species tree generated from SVD Quartets using the whole genome sequences. Colors at the tips are based on country of origin. Branches with mixed colors indicate a clade that contains samples with mixed country of origin. For example, the ancestral branch of UCBSV TZ Tan 23 KR108839 and UCBSV UG MI B3 FJ039520 is colored red and orange to indicate a clade with sampled with mixed country of origin.

### Comparison of Gene Trees to Species Tree

Clades A and B, which partition the CBSV isolates into two groups, are consistently present with high support in all genes except HAM1 and CP (Table 5). Clades D and G, which each consist of a pair of UCBSV isolates, have high support across all genes, while clades C and E have relatively high support across a majority of genes. Clade F is strongly supported by the CI gene, which is relatively long, but is not found in the phylogenetic tree estimated for any of the other genes.

The whole genome concatenated analysis using MrBayes shows strong support (posterior probability 1.0) for all clades (Table 5). However, this analysis does not take into account the possibility of variation in the evolutionary processes across the individual genes. The SVDQ analysis, on the other hand, uses a coalescentGbased method to estimate the overall species tree, and properly accounts for variation in the evolutionary history for each gene. In viewing the bootstrap support values for each of the clades from the SVDQ analysis, we see that the level of support for each clade across the genome is more accurately represented by the corresponding bootstrap proportion. For example, clade F, which was found only in the phylogeny of the CI gene, shows a bootstrap proportion of 0.44 for the SVDQ analysis (as compared to 1.0 for the MrBayes concatenated analysis) (Table 5). Similarly, the SVDQ analysis gives a bootstrap proportion of 0.87 for clade E, which showed posterior probabilities below 0.8 for 3 of the 10 genes, as compared to a posterior probability of 1.0 for the concatenated analysis with MrBayes. All other clades are supported with bootstrap values of 1.0, consistent with the MrBayes analysis.

### Sliding Window SVD Score

The SVD Score Sliding Window analysis (Fig. 4) shows several interesting patterns. First, note that the gene boundaries track well with shifts in the magnitude of the SVD Score, indicating that individual genes are subject to specific evolutionary processes that vary from gene to gene. In particular, several genes show strong support for the primary CBSV/UCBSV split, as indicated by their low scores, while other genes show variation from this basic process, as indicated by increases in the scores. In addition, Fig. 4 shows the test statistic associated with the hypothesis test for a shift in the rate of evolution between the two groups, with ‘*’ indicating that the rate difference between the two groups is statistically significant. It is readily apparent from the graph that genes that show strong support (low SVD Score) for the primary CBSV/UCBSV split also show strong evidence for statistically significant differences in evolutionary rate. These results support the overall hypothesis that certain genes in CBSV have accelerated rates of evolution that may be contributing to the higher aggressiveness of the virus.

**Figure 4.**
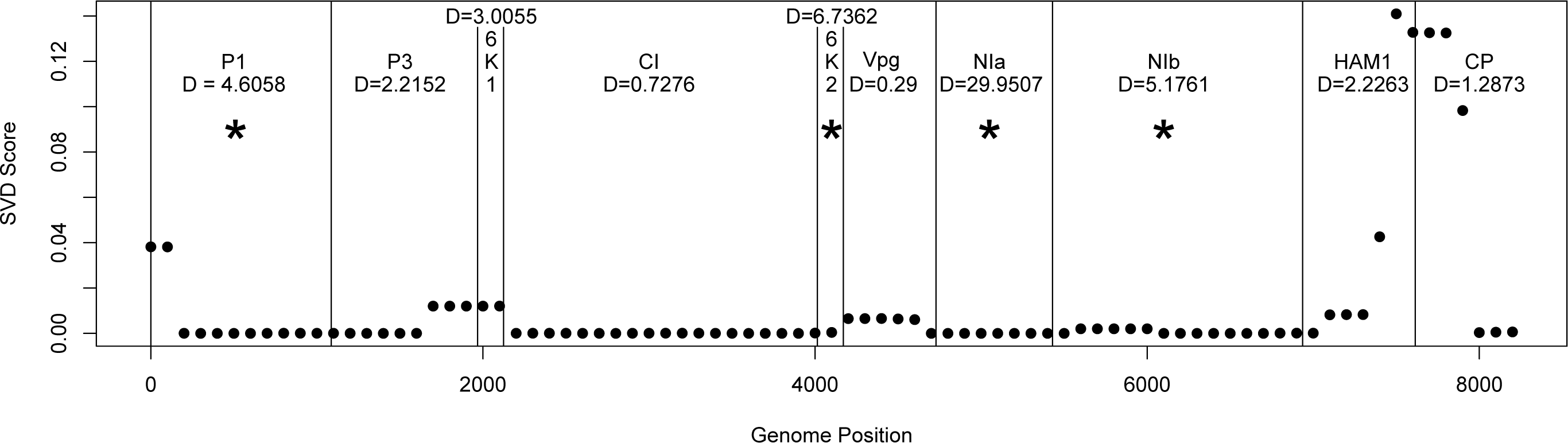
Computed SVD Score with the split defined by CBSV vs. UCBSV across the genome in windows of 500 bp, sliding in increments of 100 bp, and resulting SVD Scores plotted across the genome. Boundaries between genes are marked with vertical lines to further characterize the CBSV and UCBSV genomes. Rates of molecular evolution were estimated using CODEML implemented in PAML (Phylogenetic Analysis by Maximum Likelihood)^54^. The results are shown for each gene and D represents the difference in likelihoods from the null hypothesis (CBSV and UCBSV have equal rates) and the alternative hypothesis (CBSV and UCBSV have different rates).

## Discussion

In this study, we analyzed new and all publicly available CBSVs whole genomes to elucidate molecular mechanisms underlying the field and laboratory observations that CBSV more readily infects cassava plants and tends to display severe symptoms when compared with UCBSV. Our analyses included characterizing three new complete CBSV (2) and UCBSV (1) genomes, which were combined with the 26 previously published. Our major findings show further speciation of CBSV and UCBSV, a larger genetic landscape for CBSV, including many nonsynonymous sites, and reveal that CBSV has a faster rate of evolution compared with UCBSV (Table 4 and Fig. 4). These observations and their biological significance are discussed.

### Genes with Accelerated Rates of Evolution in CBSV

We have identified P1, 6K2, NIb and NIa as the genes with accelerated rates of evolution in CBSV. The function of P1 is as an RNA silencing suppressor (RSS), and there is also the suggestion that it may be involved in virion binding to the whitefly stylet via a “bridge” formation by a virusG encoded P1 protein for both CBSV and UCBSV. 6K2 is associated with cellular membrane and responsible for systemic infection and viral long distance movement^28^. The NIb encodes for a nuclear inclusion polymerase and the NIA for a nuclear inclusion protease^18^,^29^.

In Potyviruses generally, when NIa and VPg are associated together they are located in the cytoplasm and nucleus of infected cells. When 6K2GVPgGNIa forms a larger product, the VPg plays a role in viral RNA replication^30^. Even though VPg is not one of the genes with a higher evolution rate, both 6K2 and NIa are a part of the complex which affects replication, and this may go some way to explaining their apparent accelerated evolution rate. Is it possible that the accelerated rates of evolution for genes involved in replication could even be a response to the relatively recent interaction of the viruses and cassava? The CBSVs are not present in South America where cassava originates, so the viruses must be native to Africa. It would appear that the adaptation is still occurring and the cassava immune system does not know how to effectively fight these infections. Cassava was introduced to East Africa in the 18th century through oceanic movement. The first report of cassava brown streak disease was in 19369,^13^. There has been little opportunity for the coGevolution of the viruses and the host, therefore natural resistance would be a hard prospect. This raises the possibility of the original host of these viruses, a nonGcassava host which may be harboring the CBSVs or the most recent common ancestor of these viruses. This in turn leads us to wonder just how old these viruses and their ancestors are, if any of the CBSVs is the ancestral species, and the best way to answer such questions is to sequence more virus genomes from both cassava and nonGcassava hosts wherever they are found.

### How Can CBSV Still Function with Such a Large Genetic Landscape?

CBSV and UCBSV have different evolutionary patterns as observed by characterizing the whole genome sequences of CBSV and UCBSV separately. CBSV is genetically more diverse when compared with UCBSV, as evident by the greater amino acid usage (supplementary Fig. 1), the faster rates of evolution across the entire genome (table 4), and greater number of nonsynonymous sites across the entire genome (Fig. 2). How can CBSV still function with such a large genetic landscape? RNA viruses walk a very fine line of having the genetic arsenal to overcome the host immune system and diverging to a point that key functions of genes are lost^31^. Recent studies^32^,^33^ have shown that viruses with a large genetic landscape adapt to host changes much quicker and can overcome the host immune system faster. Viruses that occupy a large portion of the possible sequence space might be less fit but they outcompete the fitter strain when the host immune system shifts and hence these viruses have been described as adapted to “survival of the flattest”^34^,^35^. This means that a virus that covers the most sequence space will be able to adapt to host immune system faster than those with smaller spaces. Viruses that are adapted in this category (“survival of the flattest”) are going to be harder to breed resistance for because the virus has a larger ability to adapt to changes. It is clear that in our case, CBSV is the virus that has a larger sequence space (Supplemental Fig. 1) when compared to that of UCBSV, which is clearly smaller (Supplemental Fig. 2). CBSV is one of the RNA viruses that can be described as adapted to “survival of the flattest”, while UCBSV is not. Therefore, CBSV is more devastating because it has a larger genetic arsenal which it uses overcome the changes breeders are introducing into cassava.

Not only are the CBSV genomes more genetically diverse, but are also characterized by a large number of nonsynonymous changes in the genome (Fig. 2). An excess of nonsynonymous over synonymous substitutions at individual amino acid sites signifies that positive selection has affected the evolution of a protein between the extant sequences under study and their most recent common ancestor^36^. Positive selection is the process by which new advantageous genetic variants sweep a population and is the mechanism Darwin described to drive evolution. This is further evidence that might suggest that CBSV has a greater capacity to evade the cassava immune system as compared with UCBSV. CBSV had 66 sites under positive selection (Table 4) while UCBSV had none. The CBSV sites under positive selection are found not only in the regions that have gained the most attention, CP and HAM1Glike^13^, but are also found in all other genes except 6K2. This is further support for CBSV’s potential ability to outsmart the cassava immune system. Every gene in the CBSV genome (except 6K2) has sites under positive selection indicating effective RNA silencing of the virus will need to encompass many loci.

Using computational methods combined with field observations we have concluded that CBSV is more devastating than UCBSV. This assertion is also supported by two recent biological studies. The first was a test of reversion in three different cassava varieties (Albert, Kaleso and Kiroba) infected with CBSV and UCBSV. Reversion is a type of resistance mechanism whereby virusG infected plants will naturally recover from infection over time, and a proportion of their progeny from stem cuttings are virusGfree. A reversion event infers the host immune system was able to clear or restrict the virus from systemic movement. It was shown that UCBSVGinfected cassava had a higher rate of reversion when compared to plants infected with CBSV^27^ indicating that plants infected with UCBSV recovered more often than those infected with CBSV. This is another line of evidence supporting the more devastating nature of CBSV, and the possibility that cassava immune systems of the three varieties tested are struggling to resist the virus.

The second study supporting the hypothesis that CBSV is more aggressive than UCBSV analyzed virusGderived small RNAs within three cassava varieties (NASE 3, TME204 and 60444). Plants infected with viruses are known to trigger RNAi antiviral defense that can be measured by quantifying the abundance of 21G24 nucleotide (nt) segments produced by the dicer enzyme^37^. Cassava varieties were infected with either CBSV or UCBSV, NGS was used to detect virusGderived small RNAs^24^, and the 21G24 nt dicer fragments were mapped to either CBSV or UCBSV depending on which virus was used to infect the plant. The results showed that CBSV infection triggered a stronger immune response as measured by greater abundance of virus derived small RNA fragments across the entire CBSV genome compared with UCBSV. In addition, across all three genotypes they observed that cassava grafted with CBSVGinfected buds showed more severe symptoms compared to UCBSVGinfected plants^24^. This is further evidence that CBSV is a more aggressive virus and breeding for resistance to CBSV and UCBSV will require different experimental approaches.

### Implications of the Species Tree for CBSV and UCBSV

We have produced the first species tree estimate of the CBSD causal virus species using whole genome sequences and the coalescentGbased SVD Quartets species tree estimation algorithm. Differences in the evolutionary history of the two viruses are seen in the branching patterns in Figure 3. CBSV has diverged into two main clades A and B, while UCBSV has several wellG supported clades but the backbone is still unresolved, indicating more sampling is needed to fully understand the diversity and evolutionary history of UCBSV. The species tree (Fig. 3) is similar to the concatenated whole gene tree reported in Ndunguru et al.^19^, except addition of the clade labeled “G”, and lack of support for clades E and F in the UCBSV species. It is wellGdocumented that concatenating genes without using the coalescentGbased models can produce misleading results^38^,^39^. In our case, only CI supports clade F, and it is also the longest gene (1,883 bp), and therefore may swamp the signal of the other genes. The whole genome concatenation analysis recovers clade F with a posterior probability of 1.00 (Table 4). With regard to clade E, the SVDQ tree was more reflective of the individual gene tree signal by producing a bootstrap value of 0.87 versus 1.00 for the whole genome concatenated tree (Table 4). These results suggest that the estimated topology in the UCBSV species may be further refined as more samples are added.

Our integrative approach of species tree estimation coupled with analyzing rates of evolution has lead to a new framework for CBSV and UCBSV, which includes analyzing and treating these two groups of viruses as separate species. Multiple putative species of both CBSV and UCBV have been identified which means cassava needs to be resistant to the virus species that are prevalent in farmers’ fields. We argue that this genomic diversity and faster rate of evolution for CBSV is what is causing the breeders to struggle with breeding resistant varieties and also why the diagnostic primers are not working consistently. CBSV also has more positively selected sites than UCBSV. It was first thought that CBSD was restricted to the coastal areas and below 1000 m^23^ but as more genetic data is gathered CBSV and UCBSV are found at all elevations in many ecozones throughout eastern Africa^4^,^10^,^13^,^15^,^19^,^40^. We are still in the discovery phase with CBSV and UCBSV species as there are only 29 (now with the three new included here) whole genome sequences and other new species of both viruses are likely to be discovered. As we move forward it is important to include all known samples and use appropriate species tree estimation methods such as SVDQ.

Finally, the traditional gene regions (CP and HAM1Glike) that are used to delimit species and serve as the targets for diagnostic primers do not recover the species tree (Table 4). We recommend designing new diagnostic regions for other genes that recover the species tree and also do not have an accelerated rate of molecular evolution (Figure 4), such as CI or P3 for speciesGlevel diagnoses. It is possible that the spread of CBSV and UCBSV could have been exacerbated through dissemination of infected cuttings, as virus indexing with primers targeting CP may have misleadingly returned negative results.

### Implications of the Results for Cassava Breeding

During the last three decades worldwide, agricultural production has been compromised by a series of epidemics caused by new variants of classic viruses that show new pathogenic and epidemiological properties. An important determinant of the fitness of a virus in a given host is its ability to overcome the defenses of the host. Overcoming plant resistance by changes in the pathogenicity of viral populations represents a specific and important case of emergence, with tremendous economic consequences since it jeopardizes the success and durability of resistance factors in crops as an antiGviral control strategy. In this study, we found CBSV to be more variable, to have more positively selected sites, and to evolve five times faster than UCBSV. These findings have huge implications for cassava improvement efforts in Africa where CBSV is widely present. Field and laboratory results have proven CBSV to be more virulent and more devastating than UCBSV. Knowledge of specific virus species an improved cassava variety is resistant to will determine where to screen, multiply and deploy such varieties. Cassava breeders have to take into consideration the evolutionary and biological differences between CBSV and UCBSV in the breeding programs. For example, cassava breeders can breed varieties that are resistant to CBSV that can be strategically deployed in areas where CBSV is more prevalent, and similarly for UCBSV. Furthermore, it becomes more appropriate to always screen cassava materials against CBSV as a minimum, even if UCBSV is the more prevalent virus. Such a strategy will in effect ensure durable resistance as opposed to the indiscriminate screening and distribution of the improved CBSD resistant cassava varieties, without knowledge of the virus species in the area.

## Methods

### Field Plant Sample Collection

Farmers’ fields in Uganda with cassava plants 3G6 months old were surveyed for CBSD in 20 districts. In each field, cassava plants were visually assessed to confirm typical CBSD symptoms on leaves and stems. CBSD leaf symptom severity was scored on a 1G5 scale^41^,^42^; 1 = no visible symptoms, 2 = mild vein yellowing or chlorotic blotches on some leaves, 3 = pronounced/extensive vein yellowing or chlorotic blotches on leaves, but no lesions or streaks on stems, 4 = pronounced/extensive vein yellowing or chlorotic blotches on leaves and mild lesions or streaks on stems, 5 = pronounced/extensive vein yellowing or chlorotic blotches on leaves and severe lesions or streaks on stems, defoliation and dieback. CBSD symptoms were also categorized based on distribution of leaf chlorosis and stem lesions on the plant; systemic and on the whole plant (SW), systemic on leaf or stem parts but localized (SL), only on lower leaves (LL). On selected symptomatic plants, portions of the third fully expanded leaf on a shoot were picked as samples, airGdried by pressing between sheets of newsprint and stored pending RNA extraction.

### RNA Extraction

About 0.25 g cassava leaf samples were frozen in liquid nitrogen, then ground using a mortar and pestle. 2 ml CTAB lysis buffer (2% CTAB; 100 mM Tris–HCl, pH 8.0; 20 mM EDTA; 1.4 M 134 NaCl; 1% sodium sulphite; 2% PVP) was added and samples homogenized. The 1 ml of the homogenate was incubated at 65°C for 15 min, an equal volume of chloroform: isoamyl alcohol (24:1) was added, and the sample was centrifuged for 10 min at approximately 14,500rpm. 800¼l of the aqueous layer was transferred to a new tube with an equal volume of 4 M LiCl and incubated at G20°C for 2 hrs. The samples were centrifuged for 25 min at 14,500 rpm and the supernatant was poured off. The pelleted RNA was reGsuspended in 200 ¼l TE buffer containing 1% SDS, 100 ¼l of 5M NaCl. 300 ¼l of iceGcold isopropanol were added and incubated at G20°C for 30 min. The sample was centrifuged at 13,000 rpm for 10 min and the aqueous layer was decanted and RNA pellets washed in 500 ¼l of 70% ethanol by centrifuging at 13,000 rpm for 5 min. The ethanol was decanted off and RNA pellet dried to remove residual ethanol. The RNA was reGsuspended in 50 ¼l nucleaseGfree water and stored at -80°C prior to testing.

### CBSV and UCBSV Detection by RTKPCR

All samples were tested for presence of CBSV and UCBSV by a twoGstep RTGPCR assay^43^. The PCR mixture consisted of 16.0 ¼l nuclease free water, 2.5 ¼l PCR buffer, 2.5 ¼l MgCl2 (2.5 mM), 0.5¼l dNTPs (10 mM), 1.0 ¼l of each primer (10mM) [forward CBSDDF2 5’G GCTMGAAATGCYGGRTAYACAAG3’ and reverse CBSDDR 5’GGGATATGGAGAAAGRKCTCCG3’], 0.5 ¼l Taq DNA polymerase and 1.0 ¼l of cDNA. The PCR thermo profile consisted of: 94oC for 2 min followed by 35cycles of 94oC (30 s), 51oC (30 s) and 72oC (30 s) for denaturation, annealing and extension, respectively. PCR products were analysed by electrophoresis in a x1 TAE buffer on a 1.2% agarose gel, stained with ethidium bromide, visualized under UV light and photographed using a digital camera.

### Sample Selection for Sequencing

From the data obtained in the diagnostic tests, samples for sequencing were selected to represent different geographical regions, symptom types and severities. Three samples that tested positive for either CBSV (2) or UCBSV (1) were selected for this study. The two samples for which presence of CBSV was confirmed (U1 and U4) had been collected from different farmers’ fields in Mukono district, central Uganda. The sample with UCBSV (U8) selected for further analysis originated from a field in Mayuge district, eastern Uganda.

### Generation of the Transcriptomes

The three samples were transported to the laboratory and extracted as detailed above. Total RNA was blotted on to FTA cards and later extracted using methods previously described^44^. Total RNA from each sample was sent to the Australian Genome Research Facility (AGRF) for library preparation and barcoding before 100 bp pairedGend sequencing on an Illumina HiSeq2000.

### De novo Sequence Assembly and Mapping

For each sample, reads were first trimmed using CLC Genomics Workbench 6.5 (CLCGW) with the quality scores limit set to 0.01, maximum number of ambiguities to two and removing any reads with <30 nucleotides (nt). Contigs were assembled using the *de novo* assembly function of CLCGW with automatic word size, automatic bubble size, minimum contig length 500, mismatch cost two, insertion cost three, deletion cost three, length fraction 0.5 and similarity fraction 0.9. Contigs were sorted by length and the longest subjected to a BLAST search (blastn and blastx)^45^. In addition, reads were also imported into Geneious 6.1.646 and provided with reference sequences obtained from Genbank (KR108828 for CBSV and KR108836 for UCBSV). Mapping was performed with minimum overlap 10%, minimum overlap identity 80%, allow gaps 10% and fine tuning set to iterate up to 10 times. A consensus between the contig of interest from CLCGW and the consensus from mapping in Geneious was created in Geneious by alignment with MAFFT^47^. Open reading frames (ORFs) were predicted and annotations made using Geneious. Finalized sequences were designated as “complete” based on comparison with the reference sequences used in the mapping process, and “coding complete” if some of the 5’ or 3’ UTR was missing but the coding region was intact^48^,^49^, and entered into the European Nucleotide Archive (WEBIN ID number Hx2000053576).

### Genome Alignment and Annotation

TwentyGsix whole genomes (12 CBSV and 14 UCBSV) were downloaded from GenBank and imported into Geneious^46^, and the MAFFT plugin^47^was used to align them with the 3 new whole genome sequences obtained in this study. Nucleotide alignments were translated into protein using the translate align option in Geneious and then visually inspected for quality. Annotations were transferred to the 3 new genomes from the 26 previously published genomes using the live annotation option in Geneious.

### Characterizing the Genetic Diversity in CBSV and UCBSV Genomes

CBSV and UCBSV are distinct species (Fig. 2) therefore the genomes were treated separately in the analyses in characterizing the genomes. Characterizing the genetic diversity of CBSV and UCBSV was done using the <u>S</u>ynonymous <u>N</u>onGsynonymous <u>A</u>nalysis <u>P</u>rogram (SNAP v2.1.1) implemented in the Los Alamos National Laboratory HIVGsequence database (http://www.hiv.lanl.gov)^50^. SNAP calculates synonymous and nonGsynonymous substitution rates based on a set of codonGaligned nucleotide sequences. This program is based on the simple methods for estimating the numbers of synonymous and nonsynonymous nucleotide substitutions of^51^, and incorporating a statistic developed for computing variances and covariances of dS’s and dN’s^52^. An application of the SNAP package in HIVG1 research has also been developed^53^.

### Estimating Rates of Evolution

To further characterize the CBSV and UCBSV genomes, we estimated the rates of molecular evolution using CODEML implemented in PAML (Phylogenetic Analysis by Maximum Likelihood)^54^. PAML is a package of programs for analysis of DNA or protein sequences by using maximum likelihood methods in a phylogenetic framework. The null hypothesis tested was that CBSV and UCBSV have equal rates of evolution (one omega; model = 0) while the alternative hypothesis was that CBSV and UCBSV have different rates of evolution (two omegas; model = 2). The Likelihood Ratio Test was used to test for significance. If the test statistic was greater than 3.84 (based on the ChiGsquared distribution and one degree of freedom) we then rejected the null hypothesis that the rates between CBSV and UCBSV are equal. Initial analyses were carried out for the entire genome and showed that CBSV has a higher rate of evolution (Table 4). To identify which gene or genes were contributing to the faster rate of evolution we repeated the analysis separately for each individual gene.

### Testing for Positive Selection

Sites under positive selection were identified using SLAC^55^ implemented on the http://www.datamonkey.org web server^56^. The settings used to run SLAC were as follows: the best fitting model (GTR) was specified global dN/dS value was estimated and the significance level was set to 0.01.

### Gene Tree Estimation

Individual gene trees were estimated using MrBayes 3.2.157 run in parallel on Magnus (Pawsey Supercomputing Centre, Perth, Western Australia) utilizing the BEAGLE library^58^. MrBayes 3.2.1 was run utilizing 4 chains for 30 million generations and trees were sampled every 1000 generations. All runs reached a plateau in likelihood score, which was indicated by the standard deviation of split frequencies (0.0015), and the potential scale reduction factor (PSRF) was close to one, indicating that the MCMC had converged.

### Species Tree Estimation

The SVDQ method^59^ implemented in PAUP*^60^ was used to analyze the wholeGgenome data. This method enables analysis of multiGlocus data in a coalescent framework that allows for variation in the phylogenetic histories of individual genes. The method was run with all possible quartets (23,751) sampled in each of 100 bootstrap replicates, and the consensus across all bootstrap replicates was used as the estimate of the species tree. Bootstrap support values for each node were used to quantify uncertainty in the species tree estimate. The entire analysis took approximately 2.5 minutes on a MacBook Pro running OSX 10.11.2 with a 2.2 GHz Intel Core i7 processor.

### Comparison of Gene Trees to Species Tree

We compared the singleGgene phylogenies constructed using MrBayes with the overall species tree phylogeny estimated using SVDQ and the concatenated phylogeny estimated by MrBayes. For each tree, we evaluated presence or absence of the clades identified by Ndunguru et al.^19^ labeled AGF in Figure 3. We identified an additional clade (clade G, Fig. 3) that we noticed to be consistently present across genes and methods. For each of these clades present in a particular tree, we recorded the posterior probability (for trees constructed by MrBayes) or the bootstrap proportion (for the tree estimated by SVDQ) in Table 5.

### Sliding Window SVD Score

The SVD Score^61^ was used to quantify support for two viral clades for portions of the genome in a sliding window analysis. Briefly, the SVD Score measures the extent to which the data support a phylogenetic “split” – a division of the taxa into two groups with specified group membership. Low values of the SVD Score indicate strong support for the split of interest, while larger values indicate *either* a lack of support for the split or a shift in the underlying evolutionary process (see Allman et al. (2016) for details and examples). We computed the SVD Score with the split defined by CBSV vs. UCBSV across the genome in windows of 500 bp, sliding in increments of 100 bp, and plotted the resulting SVD Scores across the genome, with boundaries between genes marked with vertical lines. The computations took less than one minute on a MacBook Pro running OSX 10.11.2 with a 2.2 GHz Intel Core i7 processor.

## Acknowledgments

This work was supported by the Bill and Melinda Gates Foundation Grant no. 51466 “Regional Cassava Virus Diseases Diagnostic Project” awarded to Mikocheni Agricultural Research Institute, Tanzania, and a subGgrant to National Agricultural Research Organisation (Uganda). Computational resources provided by the Pawsey Supercomputing Centre with funding from the Australian Government and the Government of Western Australia supported this work.

## Author contributions

Sample collection was carried out by T.A., G.O., R.N., L.Kiiza. Laboratory work was carried out by G.O, R.N., L.Kiiza. J.N. P.S. M.K. Computational analyses were conducted by T.A., L.B., L.Kubatko., M.K, and all authors contributed to the writing of the manuscript.

